# Complex symbiont-pathogen interactions inhibit intestinal repair

**DOI:** 10.1101/746305

**Authors:** David Fast, Kristina Petkau, Meghan Ferguson, Minjeong Shin, Anthony Galenza, Benjamin Kostiuk, Stefan Pukatzki, Edan Foley

**Affiliations:** Department of Medical Microbiology and Immunology, Faculty of Medicine and Dentistry, University of Alberta, Edmonton, AB T6G 2S2, Canada; Department of Immunology & Microbiology, University of Colorado School of Medicine, Aurora, CO 80045

## Abstract

Pathogen-mediated damage to the intestinal epithelium activates compensatory growth and differentiation repair programs in progenitor cells. Accelerated progenitor growth replenishes damaged tissue and maintains barrier integrity. Despite the importance of epithelial renewal to intestinal homeostasis, we know little about the effects of pathogen-commensal interactions on progenitor growth. We found that the enteric pathogen *Vibrio cholerae*, blocks critical growth and differentiation pathways in *Drosophila* progenitors despite extensive damage to the epithelial tissue. We showed that inhibition of epithelial repair requires interactions of the *Vibrio cholerae* type six secretion system with a complex community of symbiotic bacteria, and that elimination of the gut microbiome is sufficient to restore homeostatic growth in infected intestines. Together, this work highlights the importance of pathogen-symbiont interactions on intestinal immune responses and outlines a previously undescribed impact of the type six secretion system on pathogenesis.

## INTRODUCTION

The digestive tract is inhabited by a dense polymicrobial community that is important for many aspects of host biology. For instance, these microbial communities induce the differentiation of immune cells, aid in the development of lymphoid tissues, and evoke specific transcriptional responses along the gut (Bouskra et al., 2008; Ivanov et al., 2008; Sommer et al., 2015). Although our understanding of the effects of the microbiome have steadily advanced, comparatively little is known about how interactions between bacteria influence the host. Because of the gut’s physiological similarity to mammals, and its simple microbiome the intestine of *Drosophila melanogaster* is a commonly used model to study host-microbe interactions. (Broderick and Lemaitre, 2012; Miguel-Aliaga et al., 2018). As the fly microbiome is cultivable there are simple protocols that allow for the generation of gnotobiotic flies that contain a defined consortium of bacteria (Douglas, 2018; Koyle et al., 2016; Ma et al., 2015). Therefore, it is possible to measure how simple interactions between two bacterial species or high-order complex interactions of more than two species impact the host (Gould et al., 2018).

To manage the intestinal microbiota, mammals and insects integrates physical, chemical, and immune defenses with homeostatic epithelial renewal to restrict the growth and dissemination of intestinal microbes, and to maintain barrier integrity. In *Drosophila*, enteric bacteria promote the synthesis of bactericidal reactive oxygen species and antimicrobial peptides that effectively prevent overgrowth of gut bacterial populations (Ha et al., 2005; Ryu et al., 2006; Tzou et al., 2000; Zaidman-Rémy et al., 2006). Damage to the gut epithelium by intestinal pathogens, or reactive oxygen species, engages reparative growth programs in intestinal progenitor cells (IPCs) that restore barrier integrity (Amcheslavsky et al., 2009; Buchon et al., 2009a; Jiang et al., 2009). Typically, infection stimulates IPC proliferation via the activation of the Epidermal Growth Factor (EGF) and Janus Kinase/Signal Transducer and Activator of Transcription (JAK/STAT) pathways (Buchon et al., 2009b, 2010; Cronin et al., 2009; Jiang et al., 2009, 2011). Immune effectors and regenerative proliferation are essential immune responses to pathogenic microbes (Miguel-Aliaga et al., 2018). However, it is important to consider how symbiotic bacteria influence the host defense response to pathogenic bacteria. For example, susceptibility to *Clostridium difficile* infection is associated with shifts in symbiotic bacteria diversity (Samarkos et al., 2018), and a decrease in the abundance of *Firmicutes* and *Bacteroidetes* alongside an expansion of *Enterobacteriaceae* (Peterfreund et al., 2012).

Approximately, 25 percent of sequenced Gram-negative bacteria encode a type six secretion system (T6SS), which injects toxic effectors into susceptible prey (Bingle et al., 2008; Das and Chaudhuri, 2003; Mougous et al., 2006; Pukatzki et al., 2006). T6SS-encoded effectors cover a range of biological functions that include phospholipid hydrolysis, actin-crosslinking, pore-formation, and peptidoglycan degradation (Miyata et al., 2011; Pukatzki et al., 2007; Russell et al., 2011, 2013). Together, these effectors permit T6SS-mediated attacks on eukaryotic and prokaryotic targets in a range of environments and hosts (Schwarz et al., 2010). Interactions between the T6SS and neighboring cells contribute to disease caused by several pathogenic bacteria. For example, the T6SS of *Campylobacter jejuni* is thought to interact with eukaryotic cells to support *in vivo* colonization (Lertpiriyapong et al., 2012). Alternatively, *Salmonella enterica* Serovar Typhimurium uses a T6SS to outcompete Gram-negative commensals and enhance colonization of the mouse intestine (Sana et al., 2016). In *Galleria mellonella*, the T6SS of *Acinetobacter baumannii* interacts with the microbiome to diminish host viability (Repizo et al., 2015). Thus, antagonistic interbacterial interactions mediated by the T6SS have measurable impacts on the virulence of intestinal pathogens. However, it remains unclear how interbacterial interactions of this nature, influence the host response to bacterial challenge.

Recently, the T6SS was demonstrated to contribute to the pathogenesis of *Vibrio cholerae* (*V. cholerae*) via interactions with the intestinal microbiome. In the infant mouse, oral infection with *V. cholerae* with a functional T6SS enhanced the development of diarrheal symptoms through interactions with symbiotic *E.coli* (Zhao et al., 2018). Previously, we showed that the T6SS of *V. cholerae* acts on Gram-negative symbionts in *Drosophila* to reduce host viability (Fast et al., 2018a). *Drosophila* is an established model for the characterization of *V. cholerae* pathogenesis (Blow et al., 2005). As in humans, adult flies are naturally susceptible to infection with *V. cholerae* and develop diarrhea-like symptoms upon infection (Blow et al., 2005). In this study, we used the *Drosophila – Vibrio* model to test how interactions between intestinal symbionts and *V. cholerae* influence host responses to intestinal challenge.

We found that the T6SS of *V. cholerae* disrupted intestinal homeostasis by blocking the regeneration of the gut epithelium. As part of a normal intestinal immune response, the gut epithelium is renewed via the proliferation of IPCs in response to infection (Bonfini et al., 2016; Buchon et al., 2009a, 2009b, 2010; Jiang et al., 2011). However, despite significant intestinal damage and extensive epithelial shedding, we did not detect an increase in IPC proliferation in guts infected with *V. cholerae* with a T6SS. Instead, we showed that the T6SS impairs growth and differentiation signals required for epithelial renewal. Strikingly, T6SS-dependent arrest of epithelial repair was the result of interactions between the microbiome and the T6SS, as ablation of the microbiome restored epithelial regeneration in response to *V. cholerae.* Furthermore, this inhibition of renewal was not the result of a bilateral interaction between *V. cholerae* and a single symbiotic species, but instead required interactions between *V. cholerae* and a multi-species consortium of intestinal symbionts. In particular, we found that interactions between *V. cholerae* and a community of three common fly symbionts are sufficient to inhibit epithelial repair, demonstrating that complex symbiont-pathogen interactions have measurable impacts on defences against pathogenic bacteria. Together, the work presented here identifies an arrest of IPC proliferation that requires interactions between the T6SS of *V. cholerae* and the intestinal microbiome.

## RESULTS

### The T6SS promotes epithelial shedding

In *Drosophila*, enteric infection results in the delamination and expulsion of damaged epithelial cells (Buchon et al., 2010; Zhai et al., 2018). To test the effect of the T6SS on epithelial delamination, we measured epithelial shedding in the guts of adult *CB>mCD8::GFP* flies infected with wildtype *V. cholerae* (C6706) or an isogenic C6706Δ*vasK* mutant, that carries an in-frame deletion in the essential T6SS gene that encodes the VasK protein (Pukatzki et al., 2006). In mock-infected, control flies, we observed few delaminating cells in the posterior midgut (Fig. 1Aa-c). In these flies, we mostly detected instances of one or two delaminating cells per gut with 90% of guts containing ten or fewer shedding cells (Fig. 1B). Infection with C6706Δ*vasK* promoted a modest increase in shedding. Specifically, we observed clusters of GFP-positive cells that typically contained fewer than ten cells per cluster, with 40% of guts containing more than ten shedding cells (Fig 1Ad-f, Fig. 1B). Infection with C6706 caused a more severe delamination phenotype that was readily visible throughout the posterior midgut (Fig. 1Ag-i). In this challenge, infected guts had multiple patches of large numbers of delaminating cells. For example, whereas 5% of samples infected with C6706Δ*vasK* had greater than 20 shedding cells in the posterior midgut, 45% of all samples infected with C6706 contained 20 or more shedding cells per area imaged in the posterior midgut (Fig. 1B). Additionally, in 10% of infected samples, challenge with C6706 caused greater than 40 shedding cells per posterior midgut, a phenotype that was absent from intestines infected with C6706Δ*vasK* (Fig. 1B). Comparisons between the treatment groups confirmed that infection with C6706 not only greatly increased the number of shedding cells per area relative to unchallenged guts (P = 4.0×10^−6^), but also increased the number of shedding cells compared to C6706Δ*vasK* (P=0.007, Fig. 1C). Together, these data demonstrate that the *V. cholerae* T6SS significantly enhances epithelial shedding in a *Drosophila* host.

**Figure 1.**
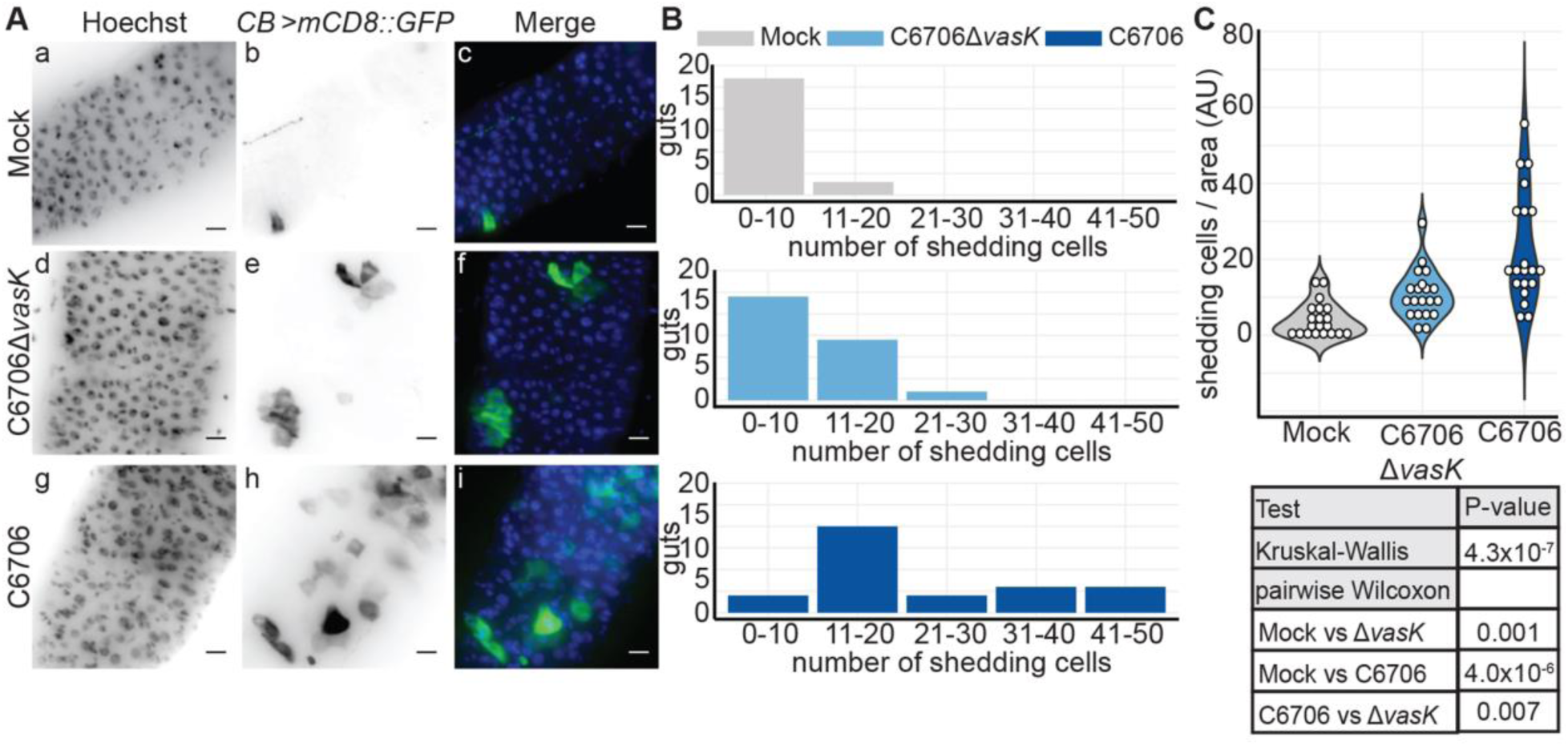
The T6SS promotes epithelial shedding. **(A)** Immunofluorescence of the posterior midgut of *CB>mCD8::GFP* flies mock infected or infected with C6706Δ*vasK,* or C6706. Scale bars are 10μm. **(B)** Histogram of the number of shedding cells in the posterior midguts from **(A)**. **(C)** Quantification of shedding cells per unit surface area from **(A)**. Each dot represents a measurement from a single fly gut.

### Disrupted intestinal homeostasis in response to the T6SS

In *Drosophila melanogaster*, intestinal damage and epithelial shedding promotes compensatory growth of IPCs to maintain the epithelial barrier (Bonfini et al., 2016). As there was extensive T6SS-dependent sloughing of epithelial cells, we tested if the T6SS promotes homeostatic growth of IPCs. To address this, we used the *esg^ts^>GFP* fly line to visualize GFP-positive IPCs in sagittal sections prepared from the posterior midguts of flies infected with C6706 or C6706Δ*vasK.* The midguts of control flies had a clear intestinal lumen surrounded by an intact epithelium (Fig. 2Aa-d). Consistent with Fig. 1, infection with C6706Δ*vasK* stimulated a modest shedding of cellular material (asterisks) into the intestinal lumen without an apparent loss of barrier integrity (Fig. 2Ae-h). Challenge with C6706 once again promoted an extensive shedding of epithelial cells and cellular debris into the lumen (Fig. 2Ae-h), as well as the appearance of numerous breaks along the basement membrane (arrowheads), suggesting pathogen-dependent damage to the epithelial barrier.

As we observed epithelial damage and shedding cells in *V. cholerae-*infected intestines, we determined if *V. cholerae* promoted compensatory growth by IPCs. In mock-infected flies, we observed the regular distribution of small GFP-positive IPCs along the basement membrane of the midgut (Fig 2Ba-d). Infection with C6706Δ*vasK* caused an accumulation of GFP-positive IPCs, consistent with enhanced epithelial renewal in response to infection (Fig. 2B e-h). In contrast, despite extensive shedding of cellular material (Fig 1) and obvious epithelial damage (Fig. 2A), guts challenged with C6706 did not appear to have elevated numbers of IPCs (Fig. 2B i-l). Instead, these guts had a limited number of basal GFP-positive cells (Fig 2Ca-d), despite an immediate proximity to lumenal bacteria (dotted outline). Taken together, these results suggest that C6706Δ*vasK* provokes a conventional intestinal immune response to pathogenic bacteria. In contrast, we did not observe signs of epithelial renewal in flies infected with C6706, despite widespread intestinal damage, raising the possibility that the *V. cholerae* T6SS uncouples epithelial shedding from intestinal regeneration.

**Figure 2.**
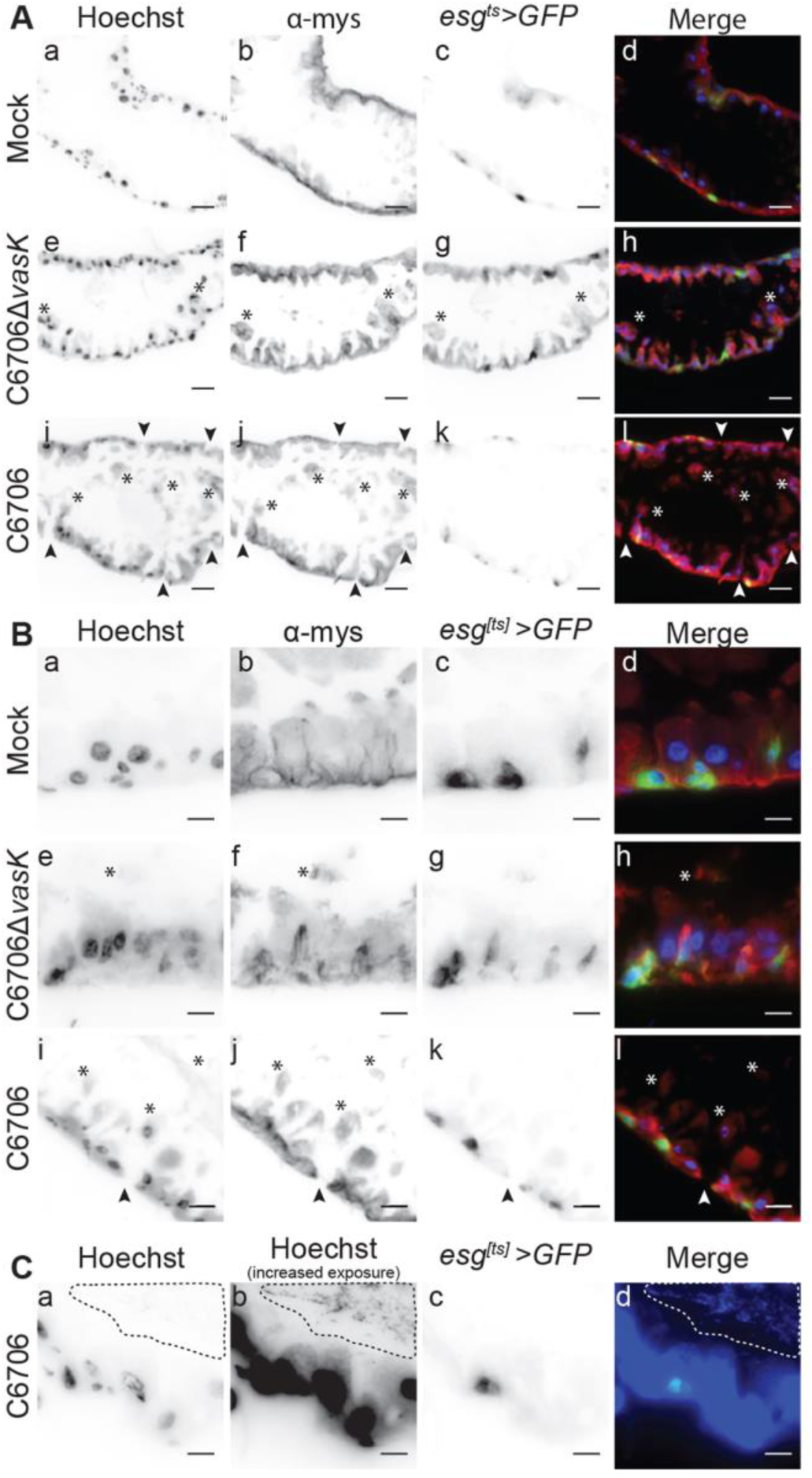
Disrupted intestinal homeostasis in response to the T6SS. **(A-C)** Immunofluorescence of sagittal sections prepared from the posterior midgut of *esg^ts^>GFP* flies mock infected or infected with C6706Δ*vasK,* or C6706. Arrowheads indicate damage to the intestinal epithelium and asterisks denote cellular matter in the lumen. **(C)** Visualization of intestinal bacteria via increased exposure of Hoechst stain. The dotted line circles bacteria in the lumen. Scale bars are **(A)** 25μm and **(B & C)** 10μm.

### The T6SS modifies IPC transcriptional responses to *V. cholerae*

To characterize effects of the T6SS on epithelial renewal, we performed RNA sequencing (RNA-seq) analysis on the intestinal response to infection with C6706 (Sup Fig. 1). We found that the host response to C6706 is characterized by the activation of antibacterial defenses, re-programming of metabolic pathways, and the expression of a large cohort of genes required for the generation and assembly of mature ribosomes. Many of these responses match our understanding of the fly transcriptional response to pathogenic bacteria (Sup Fig. 1, 4 (Buchon et al., 2009a; Dutta et al., 2015; Troha et al., 2018). However, and in contrast to classical responses to enteric challenge, we did not detect changes in mRNA levels characteristic of JAK-STAT or EGF responses, two pathways that are intimately linked with homeostatic renewal of a damaged epithelium.

The apparent absence of homeostatic growth signals in C6706-infected intestines prompted us to directly identify the transcriptional response of IPCs to *V. cholerae* infection. For this experiment, we performed RNA-seq on IPCs purified from the guts of adult *esg*[*ts*]/*+* flies that we challenged with C6706 or C6706Δ*vasK* (Fig. 3A). As a control, we sequenced the transcriptome of IPCs from uninfected *esg*[*ts*]*/ +* flies. Principle component analysis showed that samples from uninfected flies and those from flies infected with C6706Δ*vasK* grouped relatively closely. In contrast, samples from C6706-infected flies grouped away from both uninfected and C6706Δ*vasK*-infected flies (Fig. 3B). Furthermore, differential gene expression analysis revealed minimal overlaps between C6706 and C6706Δ*vasK-*infected flies relative to uninfected controls (Fig. 3C). From there, we examined Gene Ontology (GO) term enrichment among the differentially upregulated and downregulated genes. Here, we also compared C6706Δ*vasK* to C6706 to specifically identify changes in IPC transcriptional responses to the T6SS (Fig. 3F). Of note, comparison of the transcription profile of C6706-challenged IPCs with uninfected IPCs revealed a downregulation of biological processes involved in growth and mitosis. This included a significant downregulation of processes such as cell proliferation and nuclear division (Fig. 3G).

**Figure 3.**
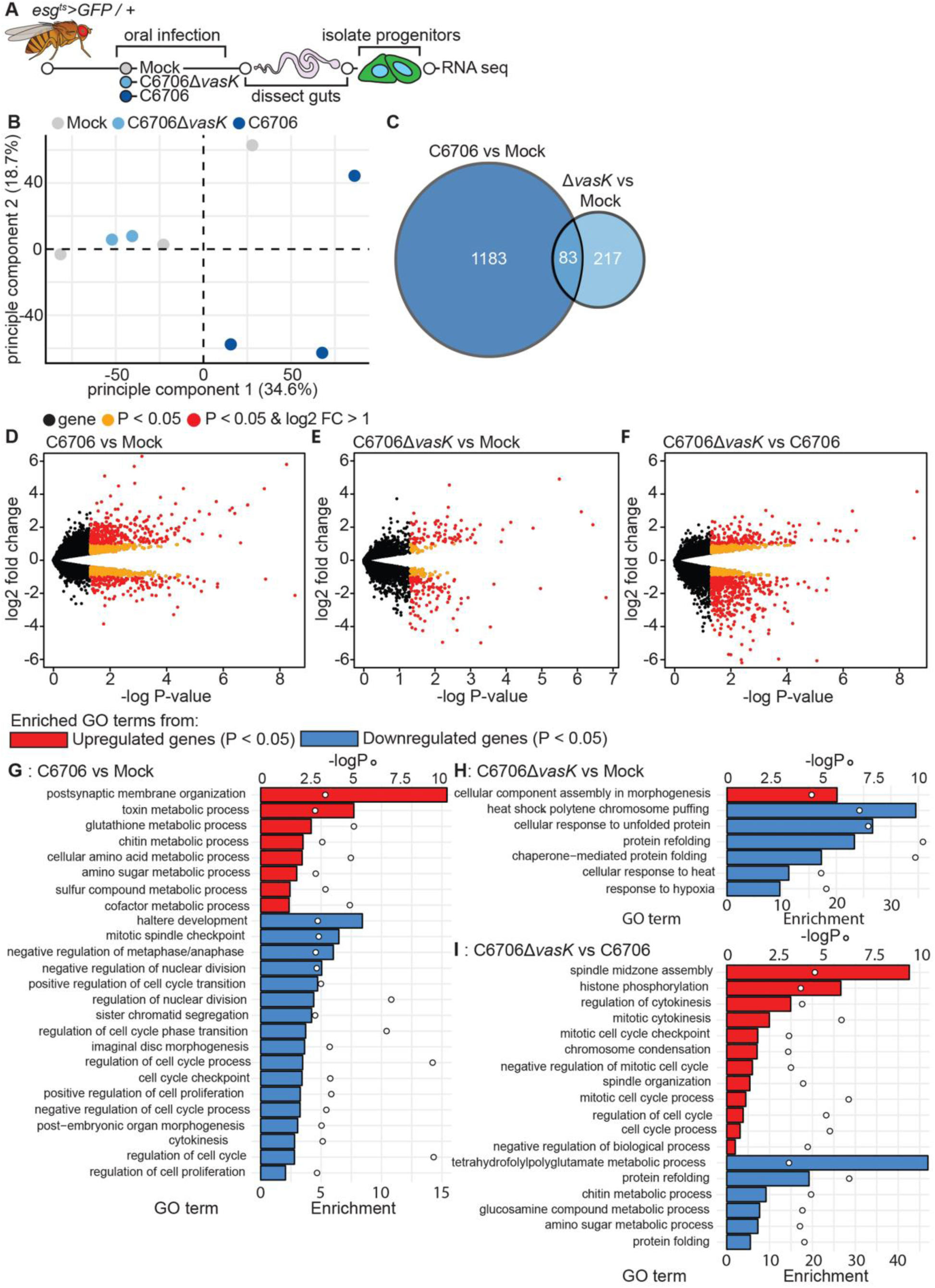
The T6SS modifies IPC transcriptional responses to *V. cholerae*. **(A)** Schematic representation of the RNA-sequencing of IPCs isolated from *V. cholerae* infected guts. **(B)** Principle component analysis from the counts per million obtained from RNA-sequencing of IPCs isolated from guts mock infected or infected with C6706 or C6706Δ*vasK.* **(C)** Venn diagram of differentially expressed genes (P<0.05) from comparisons of C6706 to Mock and C6706Δ*vasK* to Mock. **(D-F)** Volcano plots of differentially expressed genes from comparisons of **(D)** C6706 to Mock, **(E)** C6706Δ*vasK* to Mock, and **(F)** C6706Δ*vasK* to C6706. Each dot represents a single gene. Yellow indicates a P<0.05, red indicates P<0.05 and log2 fold change >1 or <-1. **(G-I)** Gene Ontology analysis from up or down regulated differently expressed genes (P<0.05) from comparisons of **(G)** C6706 to Mock, **(H)** C6706Δ*vasK* to Mock, and **(I)** C6706Δ*vasK* to C6706.

In contrast, this downregulation of growth processes was absent when we compared the transcriptional profile of C6706Δ*vasK*-infected IPCs to that of uninfected IPCs (Fig. 3H). Instead, we found a significant enrichment of mitotic processes in flies infected with C6706Δ*vasK* relative to flies challenged with C6706 (Fig. 3I). Together, these data suggest that IPCs have distinct transcriptional response to wildtype and T6SS-deficient *V. cholerae*. In particular, we found that the T6SS inhibits the expression of genes required for growth and renewal of the epithelium.

To further characterize T6SS-dependent impacts on epithelial renewal, we determined the transcriptional profile of the whole intestinal response to infection with C670*6ΔvasK* (Sup Fig. 2A). In general terms, we noticed substantial overlaps between host responses to C6706 and C6706Δ*vasK* (Sup Fig. 2B). For example, C6706Δ*vasK* caused a differential expression of genes required for the control of intestinal immunity, metabolism, and the generation of mature ribosomes (Sup Fig. 2C). However, in contrast to C6706Δ*vasK*, challenge with C6706 impacted the expression of genes required for epithelial growth and renewal, including decapentaplegic pathway elements, and core components of the cell cycle progression machinery (Sup Fig. 3) (Guo et al., 2013; Tian and Jiang, 2014; Zhou et al., 2015). Specifically, infection with C6706 resulted in a downregulation of cell cycle genes relative to challenge with C6706Δ*vasK.* These observations are in agreement with roles for the T6SS in the arrest of epithelial renewal.

### IPCs fail to mediate intestinal repair when challenged with T6SS functional *V. cholerae*

Epithelial damage activates the JAK/STAT and the EGFR pathways to stimulate epithelial repair. Consistent with this, we identified increased levels of mRNA of genes indicative of JAK/STAT and EGF pathway activation such as *argos* (*aos*) and Suppressor of cytokine signalling 36E (*Socs36E*) in IPCs from C6706Δ*vasK-*infected flies compared to those from C6706-infected counterparts (Fig. 4A). Furthermore, infection C6706Δ*vasK* led to an increase in the expression of cell cycle activators, such as the Cdc25 ortholog *string* (*stg*). We did not detect a similar engagement of repair in IPCs from flies challenged with C6706. Instead, we detected diminished levels of mRNA of a number of key signaling and regulatory components of the EGF pathway. In particular, we noted diminished expression of the EGF pathway transcription factor *pointed* (*pnt*) and the EGF receptor (*EGFR)* itself in IPCs from flies infected with C6706 compared to IPCs from uninfected controls (Fig. 4A). Similarly, we noted a reduction in the relative proportions of mRNAs that encode central components of the JAK/STAT pathway. In the JAK/STAT pathway, binding of interleukin-like ligands to the receptor Domeless (*dome*) induces signalling through the kinase Hopscotch (*hop*), and results in the transcription of *Socs36E* (Zeidler and Bausek, 2013). We observed diminished mRNA levels of all three of these signaling components in the IPCs of C6706-challenged flies relative to uninfected controls. Furthermore, we detected significant drops in mRNA that encode prominent cell cycle genes, such as *stg*, the S-phase *cyclin dependent kinase 2* (*Cdk2*), and the essential M phase cyclin *CyclinB3* (*CycB3*) in IPCs from C6706-infected flies. In summary, we detected a significant decrease in mRNA of genes in pathways responsible for epithelial renewal alongside diminished levels of cell cycle genes, indicating a downregulation of intestinal repair programs by IPCs when challenged with C6706.

**Figure 4.**
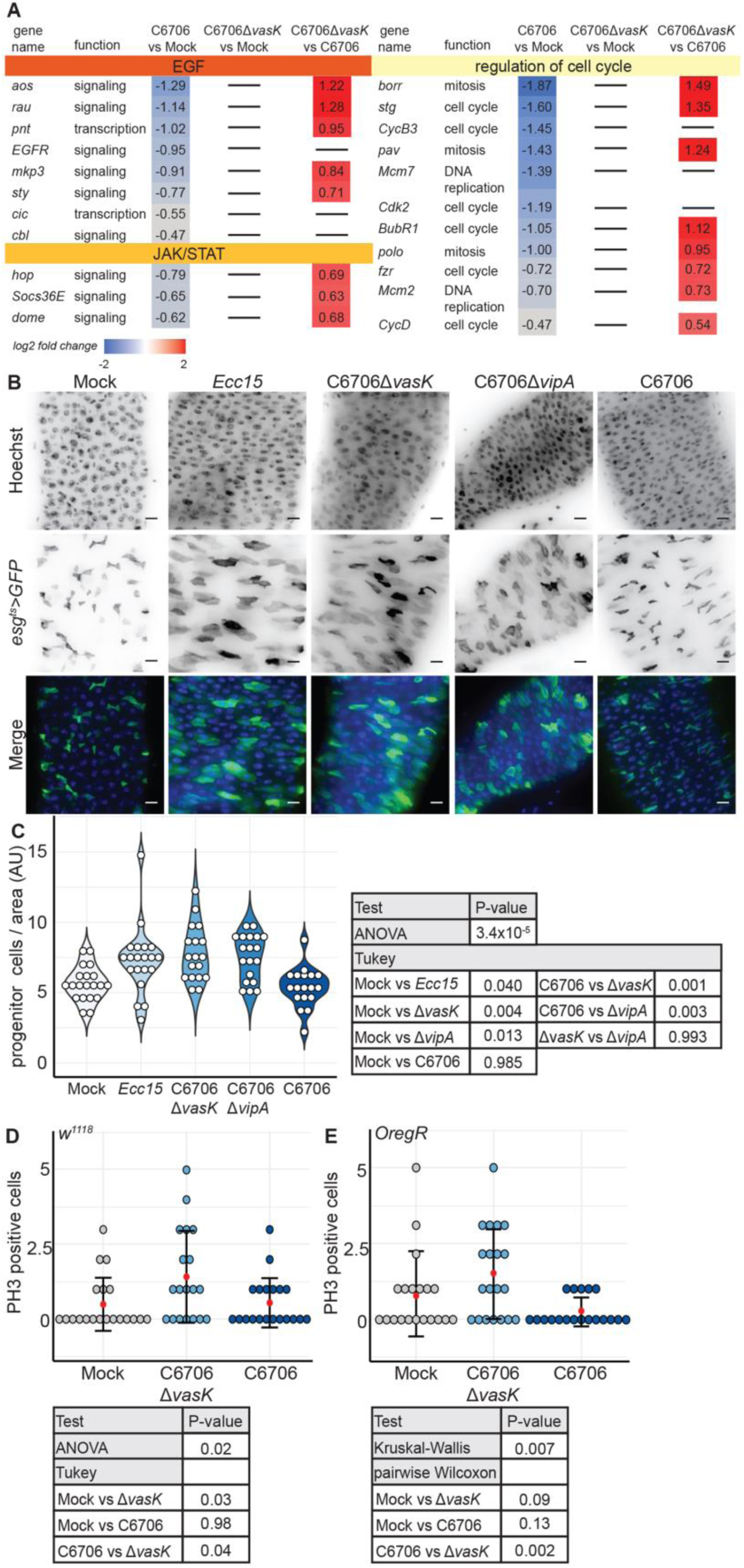
IPCs fail to mediate intestinal repair when challenged with T6SS functional *V. cholerae.* **(A)** Genes that regulate IPC growth and cell cycle from RNA-seq of IPCs of flies mock infected or infected with C6706 or C6706Δ*vasK.* **(B)** Immunofluorescence of the posterior midguts of *esg^ts^>GFP* flies mock infected or infected with *Ecc15,* C6706Δ*vasK*, C6706Δ*vipA*, or C6706. Scale bars are 10μm. **(C)** Quantification of the number of IPCs per unit surface area from **(B)**. Each dot represents a measurement from a single fly gut. **(D-E)** Quantification of the number of Ph3 positive cells in the posterior midguts of **(D)** *w^1118^* or **(E)** *OregR* flies that were mock infected or infected with C6706Δ*vasK,* or C6706.

To directly test the hypothesis that the T6SS inhibits epithelial renewal, we examined IPC growth in guts infected with C6706 or with C6706Δ*vasK* with two different functional assays. First, we quantified the number of IPCs per area in guts of infected flies as a measure of IPC expansion. As a control, we also quantified the number of IPCs in the guts of flies infected with the Gram-negative fly pathogen *Erwinia carotovora carotovora 15* (*Ecc15*), a known activator of IPC growth (Buchon et al., 2009b). In agreement with previous reports, infection with *Ecc15* promoted a significant increase in the number of IPCs per area (P=0.04, Fig. 4B, C). Similarly, guts infected with C6706Δ*vasK* had greater numbers of IPCs per area than uninfected controls (P=0.004, Fig. 4B, C). This phenotype was not specific to the *vasK* T6SS mutation, as we observed a near-identical expansion of IPCs in intestines challenged with *V. cholerae* with a null mutation in the *vipA* gene, an essential component of the T6SS outer sheath (P=0.013, Fig. 4B, C) (Zheng et al., 2011). In contrast, guts infected with C6706 had significantly fewer IPCs per area than guts infected with either C6706Δ*vasK* or C6706Δ*vipA* (P<0.001 and P<0.003 respectively, Fig. 4B, C). Furthermore, there was no difference in the number of IPCs per area between uninfected flies and those infected with C6706 (P=0.985, Fig. 4B, C), indicating a T6SS-dependent inhibition of IPC expansion. Next, we quantified mitotic PH3 positive cells in the posterior midguts of two different wildtype fly strains, *w^1118^*, and Oregon R, that we infected with C6706Δ*vasK* or C6706. In both fly backgrounds, infection with C6706Δ*vasK* prompted an increase in the number of mitotic cells in the posterior midgut. In contrast, both wildtype fly strains had significantly fewer mitotic cells in C6706-infected guts compared to C6706Δ*vask-*challenged counterparts (P=0.04 and P=0.002, Fig. 4D, E).

Collectively, these data demonstrate that the transcriptional response of IPCs to *V. cholerae* is significantly altered by the presence of a functional T6SS. This difference in response to the T6SS is highlighted by a significant downregulation of pathways critical for intestinal renewal, diminished IPC proliferation, and failed epithelial renewal.

### Impaired IPC differentiation in response to the T6SS

IPC proliferation is accompanied by signals through the Notch-Delta axis that direct the generation and differentiation of transitory enteroblasts (Micchelli and Perrimon, 2006; Ohlstein and Spradling, 2006, 2007). Our analysis of the RNA-seq data suggested T6SS-dependent effects on Notch pathway activity. For example, we detected an increase in the levels of mRNA of the Notch-response gene, *Enhancer of split* (*E(spl)*), as well as *Delta* (*Dl*) itself in IPCs from C6706Δ*vasK*-infected guts relative to C6706-infected guts (Fig. 5A). Furthermore, we noticed a suppression of *E(spl)* genes and *Dl* in IPCs from flies infected with C6706 compared to uninfected controls (Fig. 5A). As genes in the E(spl) complex are primary transcriptional targets of the Notch pathway, these data suggest a potential impairment of IPC differentiation programs by the T6SS (Bailey and Posakony, 1995).

**Figure 5.**
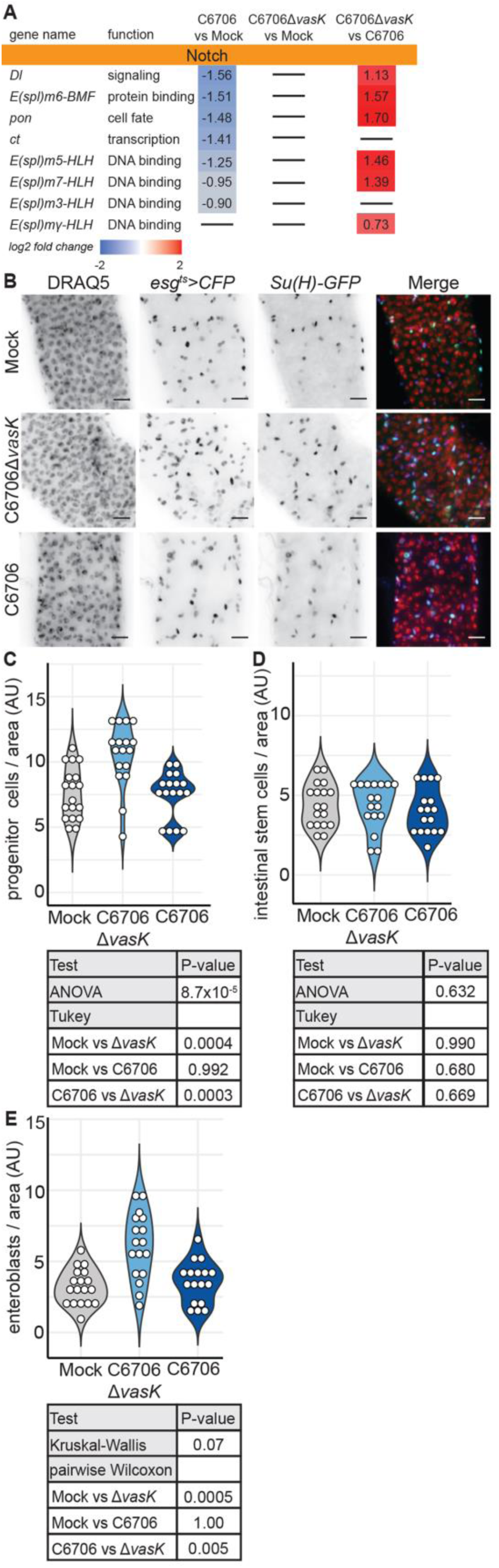
Impaired IPC differentiation in response to the T6SS. **(A)** Differentially regulated genes in the Notch signaling pathway, from RNA-sequencing of IPCs from flies mock infected or infected with C6706Δ*vasK* or C6706 **(B)** Immunofluorescence of the posterior midguts of *esg^ts^>CFP, Su(H)-GFP* flies mock infected or infected with C6706Δ*vasK*, or C6706. Scale bars are 10μm. **(C)** Quantification of the number of IPCs per unit surface area from **(B)**. Each dot represents a measurement from a single fly gut. **(D)** Quantification of the number of intestinal stem cells per unit surface area from **(B)**. **(E)** Quantification of the number of enteroblasts per unit surface area from **(B)**.

To test if IPC differentiation responds differently to the presence of a T6SS, we quantified the number of enteroblasts in the posterior midguts of flies that we infected with C6706 or C6706Δ*vasK*. In the absence of infection, we detected approximately equal numbers of intestinal stem cells (CFP-positive, GFP-negative) and enteroblasts (EB) (CFP-positive, GFP-positive) in the posterior midgut (Fig. 5B, D, E). Consistent with Figure 4, infection with C6706Δ*vasK* stimulated an expansion of IPCs (Fig. 5B, C). This expansion of IPCs was likely the result of an increased population of enteroblasts (P = 0.0004, Fig. 5E), not stem cells (Fig. 5D), consistent with the generation of undifferentiated enteroblasts required to renew the intestinal epithelium. In contrast, guts infected with C6706 contained significantly fewer IPCs per area than their C6706*ΔvasK*-infected counterparts (P = 0.0003, Fig. 5B, C). There was no difference in the number of intestinal stem cells between C6706 or C6706Δ*vasK* infected guts (Fig. 5D). Instead, there was a significant drop in the number of enteroblasts per unit area in guts challenged with C6706 relative to those infected with C6706Δ*vasK* (P= 0.005, Fig. 5B, E), indicating that the T6SS likely impairs the generation of enteroblasts.

Together, the data presented here uncover an inhibitory effect of the T6SS on epithelial renewal. We find that flies activate conventional growth and differentiation programs in response to C6706Δ*vasK.* This response is absent from intestines challenged with pathogenic *V. cholerae* with a functional T6SS. Instead, we find that despite extensive damage and increased epithelial shedding, IPCs respond with diminished levels of genes required to stimulate IPC proliferation. This change in gene expression was accompanied by diminished proliferation along with an inhibition of differentiation programs, culminating in impaired epithelial regeneration.

### IPC suppression of growth in response to the T6SS requires intestinal symbionts

T6SS effectors are toxic to eukaryotic and prokaryotic cells (Joshi et al., 2017). For example, interactions between the *V. cholerae* T6SS and eukaryotic cells have been implicated in intestinal inflammation, and recent studies have linked interactions between the T6SS and the endogenous microbiome to the virulence of *V. cholerae* (Fast et al., 2018a; Ma and Mekalanos, 2010; Zhao et al., 2018). This prompted us to ask if the IPC response to the T6SS is a function of direct interactions between the T6SS and host cells, or instead requires interactions between the T6SS and the intestinal microbiota.

To test this, we measured epithelial renewal in the guts of germ-free (GF) flies that we infected with C6706 or C6706Δ*vasK.* Similar to conventionally reared (CR) flies, which host a community of symbiotic microbes, infection of GF flies with C6706Δ*vasK* stimulated an expansion of IPCs (P= 0.00004, Fig. 6A, B). Enteric infection of GF flies with C6706 resulted in an expansion of IPCs in a manner nearly identical to that of C6706Δ*vasK*-infected intestines. Indeed, we found no significant difference in the number of IPCs per area between C6706 and C6706Δ*vasK*-infected GF flies (P = 0.658, Fig. 6A, B). These data indicate that challenges with *V. cholerae* promote epithelial renewal, and that interactions between the T6SS and the microbiota block IPC growth. To test this hypothesis, we generated germ-free flies by two different methods and measured epithelial regeneration in guts infected with C6706. Specifically, we measured the number of IPCs per area in adult germ-free flies that were generated either by administration of antibiotics to adult flies or hypochlorite dechorionation and sterilization of embryos. Here, we found that infection with C6706 promoted a significant expansion of IPCs, regardless of the method used to generate germ-free flies (P=0.0004, P=0.001, Fig. 6C), and there was no significant difference in the number of IPCs per area between antibiotic-treated or axenic flies infected with C6706 (P = 0.950, Fig. 6C). Together these results indicate that the T6SS interacts with the intestinal microbiota to impair IPC proliferation and inhibit epithelial regeneration.

**Figure 6.**
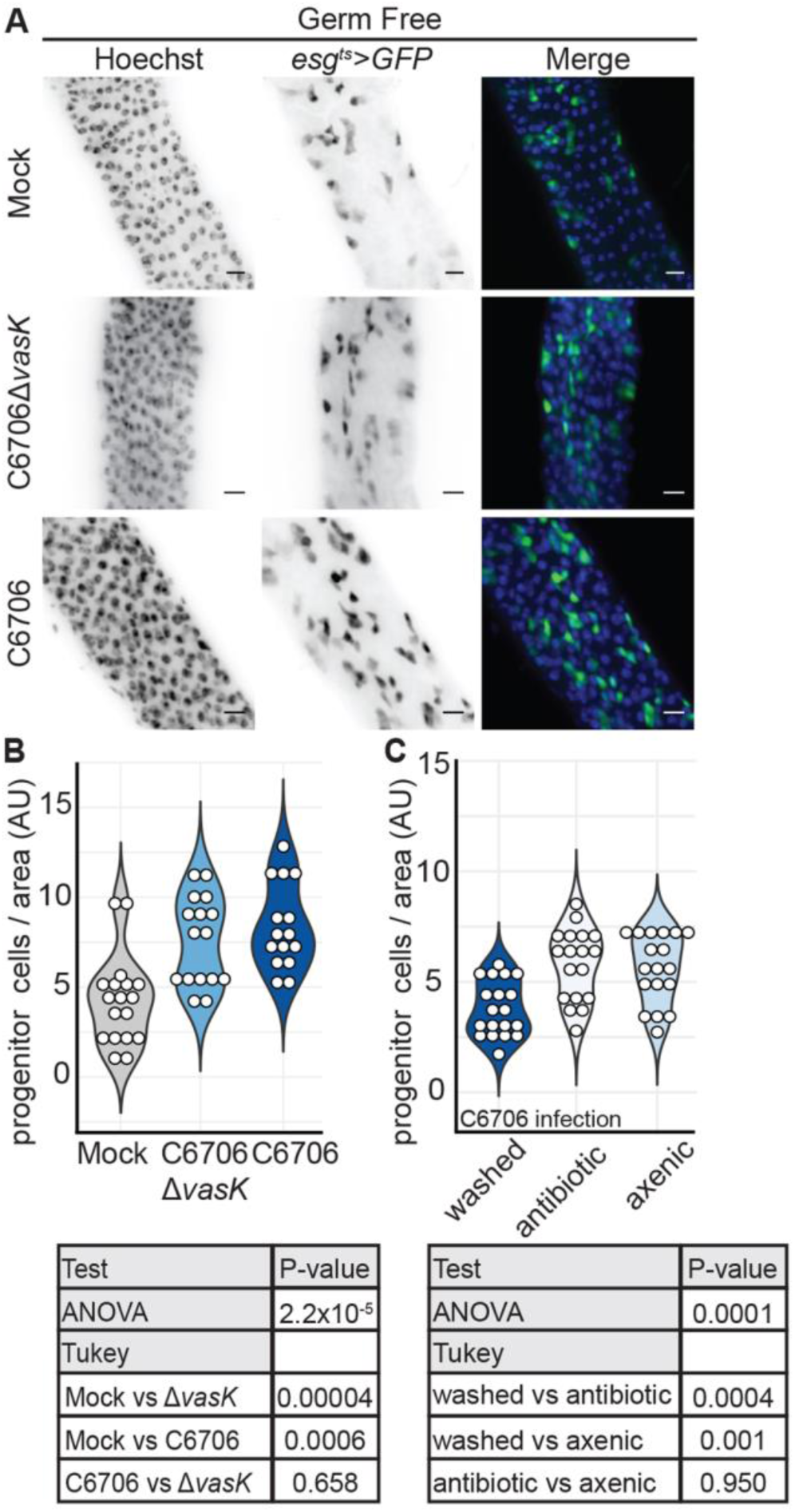
IPC suppression of growth in response to the T6SS requires intestinal symbionts. **(A)** Immunofluorescence of the posterior midguts of germ free *esg^ts^>GFP* flies mock infected or infected with C6706Δ*vasK*, or C6706. Scale bars are 10μm. **(B)** Quantification of the number of IPCs per unit surface area from **(A)**. Each dot represents a measurement from a single fly gut. **(C)** Quantification of the number of IPCs per unit surface area in *esg^ts^ >GFP* flies infected with C6706. Flies were made germ free either by the administration of antibiotics to adults (antibiotic) or by bleaching of embryos (axenic).

### T6SS suppression of epithelial renewal requires higher-order microbiome interactions

As inhibition of epithelial renewal in response to the T6SS requires gut microbes, we asked if interactions with specific members of the *Drosophila* microbiome were responsible for T6SS-mediated impairment of epithelial regeneration. We previously showed that the T6SS of *V. cholerae* targets the Gram-negative fly symbiont *Acetobacter pasteurianus* (*Ap*) for destruction, while the Gram-positive symbiont *Lactobacillus brevis* (*Lb*) is refractory to T6SS-mediated elimination (Fast et al., 2018a). As *Lb* is insensitive to the T6SS, we hypothesized that interactions between C6706 and *Lb* would fail to block epithelial repair. To test this hypothesis, we measured the number of IPCs in the guts of infected adult flies that we associated exclusively with *Lb.* For each bacterial association, we performed a parallel control infection of CR flies with the same cultures of C6706 and C6706Δ*vasK*. In each control infection, C6706Δ*vasK* promoted a regenerative response that significantly increased the number of IPCs. In contrast, challenge with C6706 consistently impaired IPC proliferation (Fig. 7A,C,D,F, G, I). We observed similar amounts of epithelial renewal in the intestines of *Lb* mono-associated flies infected with C6706 or C6706Δ*vasK* (Fig. 7B, C P=0.999), indicating that *Lb* alone does not act as an intermediary in the transmission of inhibitory-growth signals from the T6SS to the IPC. We then tested the ability of *Ap* to modify renewal. Given the sensitivity of *Ap* to T6SS-dependent killing, we expected that interactions between the T6SS and *Ap* would impair intestinal regeneration in flies challenged with C6706. However, contrary to our prediction, we did not detect a difference in the number of IPCs between *Ap*-associated guts infected with C6706 or C6706Δ*vasK* (P=0.996, Fig. 7 E. F). Instead, we found that C6706 promoted IPC proliferation when confronted with an intestine populated exclusively by *Ap*, indicating that T6SS-*Ap* interactions are not sufficient to inhibit epithelial renewal.

**Figure 7.**
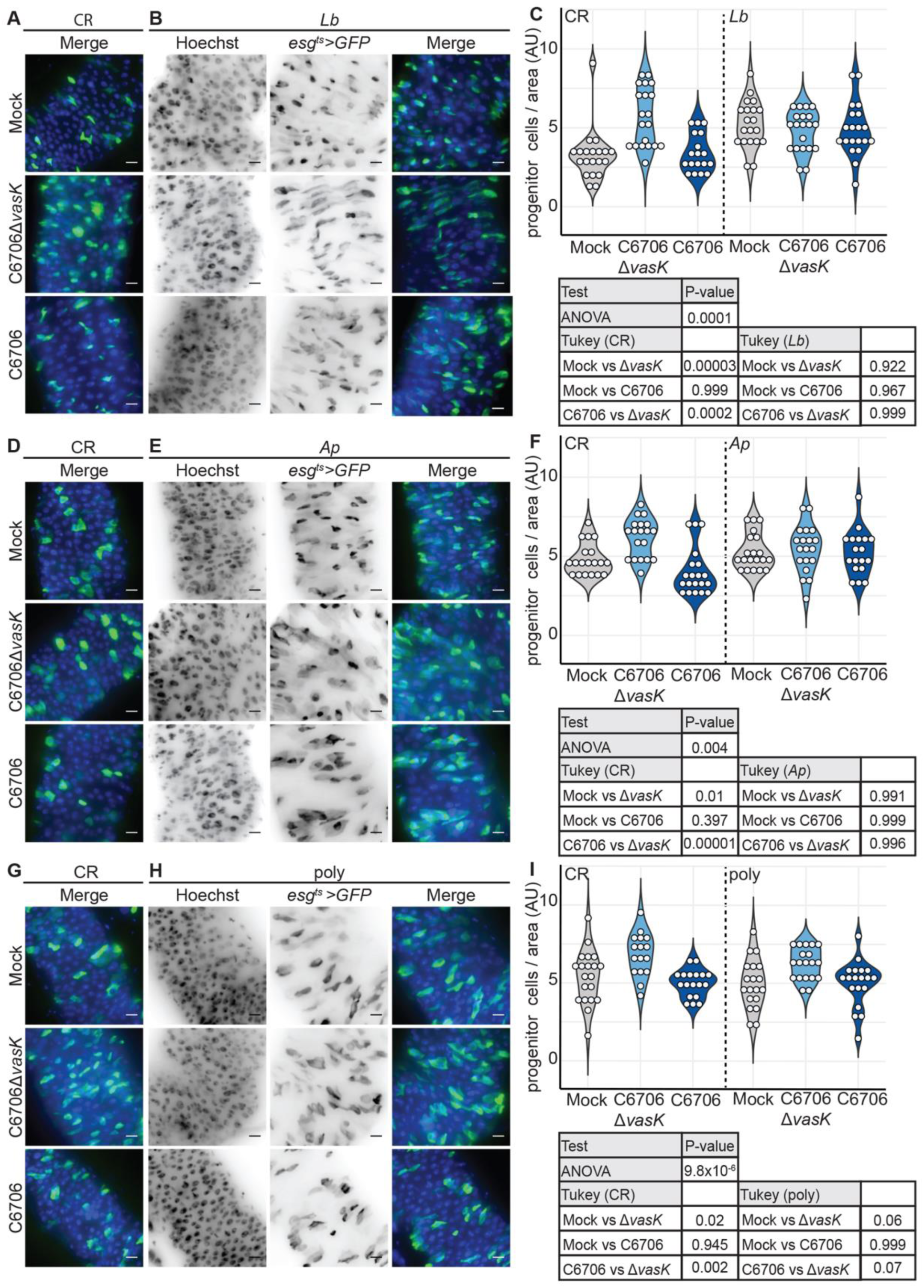
T6SS suppression of epithelial renewal requires higher-order microbiome interactions. Immunofluorescence of posterior midguts of **(A,D,G)** CR, **(B)** *Lb* mono-associated, **(E)** *Ab* mono-associated, or **(H)** poly-associated *esg^ts^ >GFP* flies mock infected or infected with C6706Δ*vasK*, or C6706. Scale bars are 10μm. Quantification of the number of IPCs per unit surface area in the guts of **(C,F,I)** CR, **(C)** *Lb* mono-associated, **(F)** *Ap* mono-associated, or **(I)** poly-associated flies. 2-3 day old virgin female flies were raised on antibiotics 5 day at 25°C to eliminate the microbiome. Germ free flies were then associated with microbial populations as indicated.

Recently, higher-order interactions among polymicrobial communities have been demonstrated to significantly influence host phenotypes in response to bacteria (Gould et al., 2018). This led us to ask if suppression of epithelial renewal by the T6SS requires a more complex community of symbiotic bacteria. To test this, we associated adult *Drosophila* with a 1:1:1 mixture of three common fly symbionts, *Ap, Lb*, and *Lactobacillus plantarum* (*Lp*), and quantified IPC numbers in the guts of flies that we infected with C6706 or C6706Δ*vasK*. Similar to what we observed in CR flies, guts infected with C6706Δ*vask* had expanded numbers of IPCs per area, indicating that poly-association with *Ap, Lb,* and *Lp,* is sufficient to reproduce physiologically relevant intestinal growth phenotypes in response to infection. In contrast, we did not see a difference in the number of IPCs between guts infected with C6706 and uninfected controls in poly-associated flies (Fig. 7H,I). Furthermore, we found an appreciable, although not statistically significant, difference in the number of IPCs between poly-associated guts infected with C6706 and C6706Δ*vasK.* These data suggest that interactions between the T6SS and individual symbiotic species are not sufficient to change IPC repair response to *V. cholerae.* Instead, impairment of epithelial renewal in response to the T6SS is the function of interactions between the T6SS and a consortium of intestinal symbionts. Together, the results presented here uncovers an inhibitory effect of the T6SS on epithelial regeneration programs, mediated by complex interactions between the T6SS and the intestinal microbiome.

## DISCUSSION

Enteric infection initiates a sequence of responses that halts the expansion and dissemination of pathogenic bacteria and mitigates intestinal damage. The renewal or turnover of the intestinal epithelium is achieved by the coordinated expulsion of damaged epithelial cells and the accelerated proliferation of IPCs. Together, these processes form an important component of the intestinal immune response (Miguel-Aliaga et al., 2018). However, it is unclear how interactions among gut-resident bacterial communities influence this response. Here, we investigated how interactions between an enteric pathogen and intestinal symbionts influence the engagement of repair programs. To explore how inter-bacterial interactions impact the intestinal response to pathogenic bacteria, we tested the effects of the T6SS, which mediates interactions between *V. cholerae* and other bacteria, on the gut transcriptional response, epithelial shedding, and IPC proliferation. We found that infection with the T6SS mutant, C6706Δ*vasK,* matched our understanding of the gut’s response to enteric infection. Specifically, challenge with C6706Δ*vasK* promoted transcription of antimicrobial peptides (Sup Fig. 4), shedding of epithelial cells, and the engagement of IPC proliferation and differentiation. These data demonstrate that infection with a T6SS deficient *V. cholerae* induces a classical immune response in the host. However, infection with C6706, which encodes a fully functional T6SS, significantly altered the host response to challenge with *V. cholerae,* indicating a previously unknown effect of the T6SS on host intestinal responses.

While infection with C6706 also promoted antimicrobial peptide transcription (Sup Fig. 4), guts infected with C6706 were phenotypically distinct from those challenged with a T6SS null mutant. In particular, C6706 promoted a more extensive shedding of epithelial cells and induced a different IPC transcriptional response. This change in IPC transcription was characterized by a downregulation of growth signals vital to the engagement of intestinal repair. Consequently, infection with C6706 effectively blocked the reparative proliferation of IPCs. Strikingly, we found that interactions between the T6SS of *V. cholerae* and the microbiome were responsible for the inhibition of intestinal regeneration. Specifically, this impaired response to bacterial challenge was the result of complex interactions that required a consortium of symbiotic bacteria, rather than a simple bilateral transaction between the pathogen and an individual symbiotic species. Together, our work details a previously unidentified consequence of infection with a bacteria with a T6SS, and sheds light on the importance of interbacterial interactions within the host.

Previously, we found that interactions between the symbiotic species *Ap* and the T6SS of *V. cholerae* combined to reduce host viability (Fast et al., 2018a). However, interactions between *Ap* and *V. cholerae* are not sufficient to impair intestinal regeneration. This suggests that the reduction in host viability and the impairment of IPC proliferation in response to the T6SS are independent consequences of intestinal challenge with *V. cholerae*. One explanation for why interactions between *Ap* and the T6SS are not sufficient to inhibit proliferation comes from the effect of *Ap* mono-association on IPCs. *Ap* promotes growth and renewal of the intestinal epithelium (Fast et al., 2018b). As it is highly unlikely that *V. cholerae* targets each *Ap* bacterium in a mono-associated gut, it is possible a portion of *Ap* promotes IPC proliferation despite the generation of putative pathogenic signals from T6SS-*Ap* interactions. This is supported by the finding that interactions between the T6SS and a multi-species community of microbes results in stunted epithelial renewal in response to *V. cholerae.* In this setting, the effects of other symbiotic species may alter or dampen the proliferative response of IPCs to *Ap,* and thereby permit inhibition of epithelial renewal in response to *V. cholera*e. The change of an effect of a single symbiotic species by the presence of other bacteria is consistent with a recent report that species diversity significantly impacts the effect of a particular symbiotic species on host physiology (Gould et al., 2018).

In *Drosophila*, challenges with large doses of *Pseudomonas entomophila* induce a translational blockade that leads to diminished repair responses in the gut (Bonfini et al., 2016; Buchon et al., 2009b). Our work matches an earlier report that showed a lack of IPC mitosis in the guts of flies infected with C6706 (Kamareddine et al., 2018). However, in contrast to *Pseudomonas entomophila*, *V. cholerae* inhibition of epithelial renewal requires interactions between the T6SS and the gut microbiota. At present, we do not understand the mechanism by which interactions between C6706 and the microbiome inhibit IPC-mediated repair. However, there are several possible explanations for this effect. The gut has co-evolved with intestinal symbionts such that the fly is sensitive to growth cues received or generated through host-microbe interactions (Broderick et al., 2014; Buchon et al., 2009b; Jones et al., 2013; Shin et al., 2011). Thus, it is possible that interactions between the microbiome and *V. cholerae* generate a different set of signals that arrest IPC proliferation, rather than stimulate division. Alternatively, interactions between symbionts and the pathogen may result in a scenario where the anti-eukaryotic function of the T6SS comes into play. For example, we measured an increase in the number of shedding epithelial cells in the guts of flies infected with C6706 (Fig. 1). Thus, it is possible that pervasive epithelial shedding is stimulated by the interaction between *V. cholerae* and the microbiome. This excess shedding may permit access of *V. cholerae* to IPCs that would otherwise be protected by the epithelium. In this context, the T6SS is capable of intoxicating eukaryotic cells with the actin crosslinker, VgrG-1. (Pukatzki et al., 2006, 2007). Together, damage induced by these eukaryotic effectors may be sufficient to halt IPC division and thereby prevent engagement of reparative programs. Future studies should consider examining the role of VgrG-1 as downstream mediators of T6SS-dependent toxicity.

The model that excessive shedding permits access of *V. cholerae* to IPCs is consistent with a recently described role for the antibacterial Immune deficiency (IMD) pathway in intestinal immunity. In the gut of *Drosophila,* IMD controls the production of antimicrobial peptides, and regulates the shedding of epithelial cells (Bosco-Drayon et al., 2012; Tzou et al., 2000; Zhai et al., 2018). Previously, we and others reported that flies with null mutations in the IMD pathway outlive wild-type flies when infected with *V. cholerae* (Fast et al., 2018a; Wang et al., 2012). This raises the possibility that secondary responses in the host contribute to the pathogenesis of *V. cholerae.* Given the recent evidence that IMD controls the sloughing of epithelial cells, it is possible that null mutations in the IMD pathway prevent excess epithelial shedding, and thereby maintain a barrier that protects IPCs from exposure to *V. cholerae.* This is further supported by our recent study which found that inhibition of IMD pathway activity exclusively in enterocytes extended the viability of flies infected with C6706 (Shin et al., 2019).

Studies in mammals, fish and insects showed that enteric microbes significantly alter the transcriptional profile of the host (Bost et al., 2018; Broderick et al., 2014; Rawls et al., 2006). Here, we identified a potent down-regulation of genes involved in homeostatic epithelial renewal in IPCs challenged with *V. cholerae*. The extent of this response was such that it covered multiple components of the EGFR, JAK/STAT, and Notch signaling pathways. Given the breadth of this response, going forward it would seem prudent to explore how this suppression of canonical growth genes is accomplished by the cell.

In summary, the work presented here demonstrates that complex interactions between intestinal symbionts and enteric invaders combine to influence critical components of the intestinal immune response. While the effects of pathogenic bacteria on epithelial repair have been described previously, our work takes in to consideration how interactions between bacterial species within a complex community structure affects this process and uncover a previously unknown effect of the T6SS. Given the diversity of intestinal microbial communities, we believe these findings represent a valuable contribution to the understanding of the effects of the microbiome on host immunity.

## METHODS

### EXPERIMENTAL MODEL AND SUBJECT DETAILS

#### Bacterial stocks

All *Drosophila* symbiotic bacterial strains were isolated from wild type lab flies in the Foley lab at the University of Alberta. *Lactobacillus plantarum KP* (DDBJ/EMBL/GenBank chromosome 1 accession CP013749 and plasmids 1-3 for accession numbers CP013750, CP013751, and CP013752, respectively), *Lactobacillus brevis EF* (DDBJ/EMBL/GeneBank accession LPXV00000000), and *Acetobacter pasteurianus AD* (DDBJ/EMBL/GeneBank accession LPWU00000000). *Lactobacillus plantarum KP, Lactobacillus brevis EF,* and *Acetobacter pasteurianus AD* have previously been described (Fast et al., 2018b; Petkau et al., 2016). *Lactobacillus plantarum* was grown in MRS broth (Sigma Lot: BCBS2861V) at 29°C for 24hours. *Lactobacillus brevis* was grown in MRS broth at 29°C for 48hours. *Acetobacter pasteurianus* was grown in MRS broth at 29°C with shaking for 48hours. *Vibrio cholerae* C6706 (a gift from John Mekalanos), C6706Δ*vasK,* and C6706Δ*vipA* have previously been described (Pukatzki et al., 2006; Zheng et al., 2011). *Vibrio* strains were grown in Lysogeny Broth (LB) (1% tryptone, 0.5% yeast extract, 0.5% NaCl) at 37°C with shaking in the presence of 100 μg/ml streptomycin. *Erwinia carotovora carotovora15* (a gift from Nicholas Buchon) was grown in LB (Difco Luria Broth Base, Miller. BD, DF0414-07-3) medium at 29°C with shaking for 24hours.

#### *Drosophila* stocks and husbandry

All fly stocks were maintained at either 18°C or 25°C on standard corn meal medium (Lakovaara, 1969). All experimental flies were adult virgin females. Fly lines used in this study were *w; upd2_CB-GAL4, UAS-mCD8:: GFP;* (a gift from Bruno Lemaitre, (Zhai et al., 2018), *w; esg-Gal4, tub-Gal80^TS^, UAS-GFP;* (referred to as *esg^ts^,* a gift from Bruce Edgar, (Micchelli and Perrimon, 2006), *w^1118^*, OregR, and *w; esg-Gal4, tub-Gal80^TS^, UAS-CFP, Su(H)-GFP;* (a gift from by Lucy O’Brien).

To make germ free flies by antibiotic treatment, freshly eclosed adult flies were raised on autoclaved standard medium supplemented with an antibiotic solution (100 g/ml ampicillin (Sigma BCBK5679V), 100 g/ml metronidazole (Sigma SLBG3633V), 50 g/ml vancomycin (Sigma 057M4022V) dissolved in 50% ethanol, and 100 g/ml neomycin (Sigma 071M0117V) dissolved in water) to eliminate the microbiome from adult flies (Ryu et al., 2008). Conventionally reared counterparts were raised on autoclaved standard cornmeal medium.

To generate axenic flies, embryos were laid on apple juice plates over a 16-h period and collected. The following steps were performed in a sterile tissue culture hood. Embryos were rinsed from the plate with sterile phosphate-buffered saline (PBS). Embryos were placed in a in a 10% solution of 7.4% sodium hypochlorite (Clorox 02408961) for 2.5 minutes, then placed into fresh 10% sodium hypochlorite solution for 2.5 minutes, and then washed with 70% ethanol for 1 minute. Embryos were then rinsed 3 times with sterile water, placed onto sterile food, and maintained at 25°C in a sterilized incubator (Koyle et al., 2016). Prior to infection or symbiont association, microbial elimination from adult flies was confirmed for every vial of axenic or germ-free flies by plating whole-fly homogenates on agar plates permissive for the growth of *Lactobacillus* and *Acetobacter*.

### METHOD DETAILS

#### Generation of gnotobiotic *Drosophila*

Virgin females were raised on antibiotic-supplemented fly food for 5 days at 25°C. On day 5 of antibiotic treatment, a fly from each vial was homogenized in MRS broth and plated on MRS and GYC agar plates to ensure eradication of the microbiome. Flies were starved in sterile empty vials for 2 h prior to bacterial association. For mono-associations, the optical density at 600 nm (OD600) of bacterial liquid cultures was measured and then the culture was spun down and resuspended in 5% sucrose in PBS to a final OD600 of 50. For poly-associations, bacterial cultures of *A. pasteurianus, L. brevis*, and *L. plantarum* were prepared to an OD600 of 50 in 5% sucrose in PBS as described above. The bacterial cultures were then mixed at a 1:1:1 ratio. For all bacterial associations, 12 flies/vial were associated with 1 ml of bacterial suspension on autoclaved cotton plugs (Fisher Scientific Canada, 14127106) in sterile fly vials. Flies were fed the bacteria-sucrose mixture for 16 h at 25°C and then flipped onto autoclaved food and raised for 5 days at 29°C. Conventionally reared control flies were given mock associations of 1 ml of 5% sucrose in sterile PBS for 16 h at 25°C. To ensure bacterial association, a sample fly from every vial was homogenized in MRS broth and plated on MRS 1 day prior to infection.

#### Immunofluorescence

Flies were washed with 95% ethanol and dissected in PBS to isolate adult intestines. Guts were fixed for 1hour at room temperature in 8% formaldehyde in PBS. Guts were rinsed in PBS for 20 minutes at room temperature and blocked overnight in PBT + 3% BSA (Sigma SLBW6769) (PBS, 0.2% Triton-X) at 4°C. Guts were stained overnight at 4°C in PBT + 3% BSA with appropriate primary antibodies, washed with PBT and stained for 1 hour at room temperature with appropriate secondary antibodies. Guts were rinsed with PBT and then stained with DNA dye for 10 minutes at room temperature. Guts were then rinsed in PBT and a final wash in PBS. Guts were mounted on slides in Fluoromount (Sigma-Aldrich F4680), and R4/R5 region of the posterior midgut was visualized. For sagittal sections, the posterior midgut was excised from dissected whole guts and imbedded in clear frozen section compound (VWR, 95057-838). Guts were cryosectioned in 10μm sections at the Alberta Diabetes Institute Histocore at the University of Alberta. All guts were visualized with a spinning disk confocal microscope (Quorum WaveFX; Quorum Technologies Inc.). Images were collected as z-slices and processed and with Fiji software to generate a single z-stacked image and measure gut area. The primary antibodies used in this study were as follows: anti-PH3 (1:1000, Millipore (Upstate), 06-570), anti-GFP (1:1000, Invitrogen, G10362), anti-myospheroid (1:100, CF.6G11 was deposited to the DSHB by Brower, D. DSHB Hybridoma Product CF.6G11). The secondary antibodies used in this study were goat anti-rabbit Alexa Fluor 488 (1:1000, Invitrogen, 1981125) and goat anti-mouse Alexa Fluor 568 (1:1000, Invitogen, 1419715). DNA stains used in this study were Hoechst 33258 (1:500, Molecular Probes Life Technologies, 02C1-2) and DRAQ5 (1:400, Invitrogen, 508DR0200G).

#### Oral infection

All infections in this study were administered orally. Virgin female flies were separated from male flies after eclosion and placed on autoclaved standard Bloomington food for 5 days at 29°C without flipping. Flies were starved 2 hours prior to infection. For *Vibrio* infections, *V. cholerae* was grown on LB plates (1% tryptone, 0.5% yeast extract, 0.5% NaCl, 1.5% agar) at 37°C in the presence of 100 μg/ml streptomycin (Sigma SLBK5521V). Colonies were suspended in LB broth and diluted to a final OD600 of 0.125. For each infection group, groups twelve flies were placed in four vials containing one third of a cotton plug soaked with 3ml of sterile LB (Mock) or with LB containing *V. cholerae*. For infection with *Erwinia, Ecc15* was grown medium at 29°C with shaking for 24hours and gathered by centrifugation. The pellet was then re-suspended in the residual LB, and 1ml of the suspension was pipetted onto a thin slice of a cotton plug at the bottom of a sterile fly vial. For all infections in this study all flies were kept on their respective infections for 24hours.

#### IPC isolation and RNA extraction

IPC isolation by fluorescence activated cell sorting (FACS) was adapted from (Dutta et al., 2013). In brief, three biological replicates consisting of 100 fly guts per replicate with the malpighian tubules and crop removed were dissected into diethyl pyrocarbonate (DEPC) PBS and placed on ice. Guts were dissociated with 1mg/ml of elastase at 27°C with gentle shaking and periodic pipetting for 1hour. IPCs were sorted based on GFP fluorescence and size with a BD FACSAria IIIu. All small GFP positive cells were collected into a tube containing DEPC PBS. Cells were pelleted at 500G for 20 minutes and then resuspended in 500μl of Trizol (ThermoFisher 155596026). Samples were stored at −80°C until all samples from each group were collected. RNA was isolated via a standard Trizol chloroform extraction. Purified RNA was sent on dry ice to the Lunenfeld-Tanenbaum Research Institute (Toronto, Canada) for library construction and sequencing. The sample quality was evaluated using Agilent Bioanalyzer 2100. TaKaRa SMART-Seq v4 Ultra Low Input RNA Kit for Sequencing was used to prepare full length cDNA. The quality and quantity of the purified cDNA was measure with Bioanalyzer and Qubit 2.0. Libraries were sequenced on the Illumina HiSeq3000 platform. For RNA-sequencing of whole guts, RNA was extracted in biological triplicate consisting of 10 dissected whole guts per replicate. RNA was purified by standard TRIZOL chloroform protocol. Purified RNA was sent on dry ice to Novogene (California, USA) for poly-A pulling, library construction and sequencing with Illumina Platform PE150 (NOVAseq 600). The sample quality was evaluated before and after library construction using an Agilent Bioanalyzer 2100.

#### Read processing, alignment, differential expression, and GO analysis

For RNAseq studies, we obtained on average 30 million reads per biological replicate. We used FASTQC (https://www.bioinformatics.babraham.ac.uk/projects/fastqc/, version 0.11.3) to evaluate the quality of raw, paired-end reads, and trimmed adaptors and reads of less than 36 base pairs in length from the raw reads using Trimmomatic (version 0.36) (Bolger et al., 2014). HISAT2 ((version 2.1.0) (Kim et al., 2015) was used to align reads to the *Drosophila* transcriptome-bdgp6 (https://ccb.jhu.edu/software/hisat2/index.shtml), and converted the resulting BAM files to SAM flies using Samtools (version 1.8) (Li et al., 2009). Converted files were counted with Rsubread (version 1.24.2) (Liao et al., 2013) and loaded into EdgeR (McCarthy et al., 2012; Robinson et al., 2010). In EdgeR, genes with counts less than 1 count per million were filtered and libraries normalized for size. Normalized libraries were used to call genes that were differentially expressed among treatments. For IPC RNA-seq, genes with P-value < 0.05 were defined as differentially expressed genes. For whole gut RNA-seq, Genes with P-value < 0.01 and FDR < 0.01 were defined as differentially expressed genes Principle component analysis was performed on normalized libraries using Factoextra (version 1.0.5) (Alboukadel and Mundt, 2017), and Gene Ontology enRIchment anaLysis and visuaLizAtion tool (GORILLA) was used to determine Gene Ontology (GO) term enrichment (Eden et al., 2009). Specifically, differentially expressed genes were compared in a two-list unraked comparison to all genes output from edgeR as a background set. Redundant GO terms were removed.

### QUANTIFICATION AND STATISTICAL ANALYSIS

#### Statistical analysis and data visualization

All graphs, plots, Venn diagrams, and GO-term lists were constructed using R (version 3.5.1) via R-studio (version 1.1.463) with ggplot2 (version 3.1.1). All figures were assembled using Adobe Illustrator. All statistical analysis was completed with R. Normality of data was determined by Bartlett test for equal variances. For normal data, one-way Analysis of Variance (ANOVA) was used to determine overall statistical difference and a Tukey’s test for Honest Significant Differences was used for multiple comparisons. For non-normal data, a Kruskal-Wallis test was used to determine overall statistical difference and pairwise Willcoxon tests with a Bonferroni correction for multiple comparisons was used for multiple comparisons.

### DATA AND CODEAVAILABILITY

#### Data availability

Gene expression data have been submitted to the NCBI GEO database (GSE136069).

## SUPPLEMENTAL INFORMATION

**Supplemental Figure 1.**
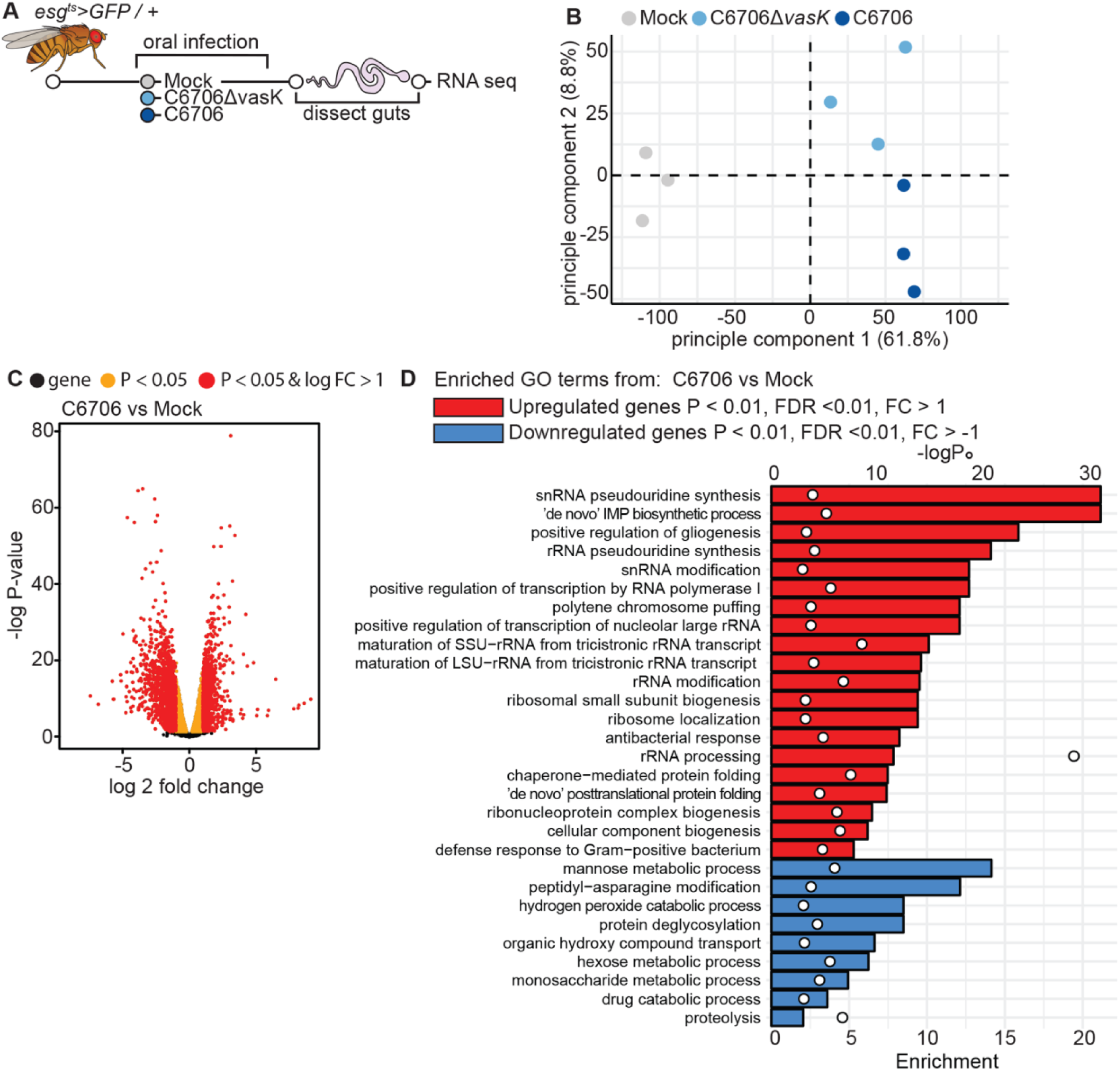
The T6SS modifies whole gut transcriptional responses to *V. cholerae.* **(A)** Schematic representation of the RNA-sequencing of *V. cholerae* infected guts. **(B)** Principle component analysis from the counts per million obtained from RNA-sequencing of guts dissected from mock infected flies or flies infected with C6706 or C6706Δ*vasK.* **(C)** Volcano plots of differentially expressed genes from comparison of C6706 to Mock. Each dot represents a single gene. Yellow indicates a P<0.05 and red indicates P<0.05 and log2 fold change >1 or <-1. **(D)** Gene Ontology (GO) analysis from the top 500 up or down regulated differently expressed genes (P<0.01, FDR<0.01, and log2 fold change >1 or < −1) from comparisons of C6706 to Mock

**Supplemental Figure 2.**
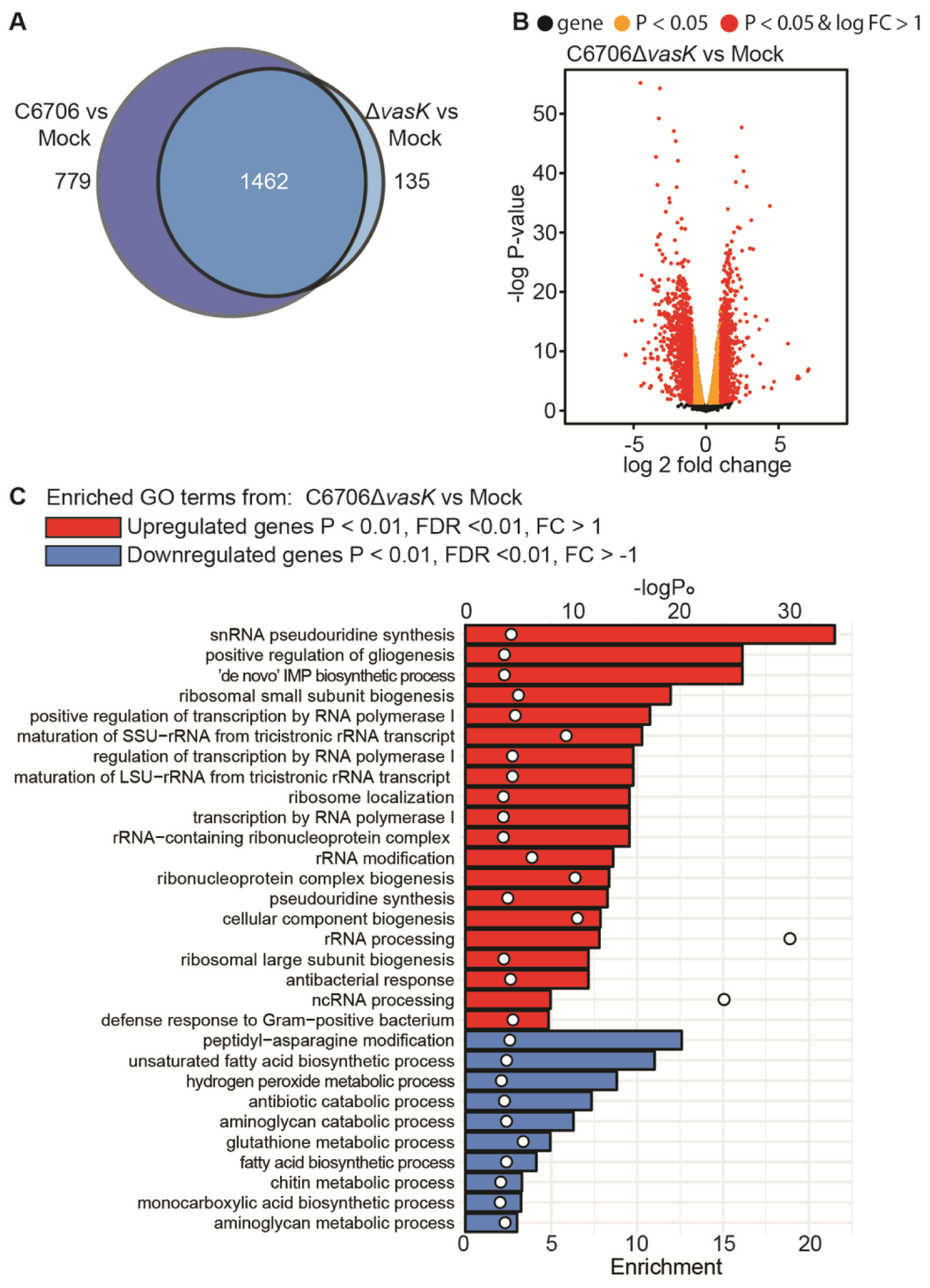
The gut transcriptional responses to C6706Δ*vasK.* **(A)** Venn diagram of differentially expressed genes (P<0.01, FDR< 0.01, and log2 fold change >1 or < −1) from comparisons of C6706 to Mock and C6706Δ*vasK* to Mock. **(B)** Volcano plot of differentially expressed genes from comparison of C6706Δ*vasK* to Mock. Each dot represents a gene. Yellow indicates a P < 0.05 and red indicates P<0.05 and log2 fold change >1 or <-1. **(C)** Gene Ontology (GO) analysis from the top 500 up or down regulated differently expressed genes (P<0.01, FDR < 0.01, and log2 fold change >1 or < −1) from comparisons of C6706Δ*vasK* to Mock.

**Supplemental Figure 3.**
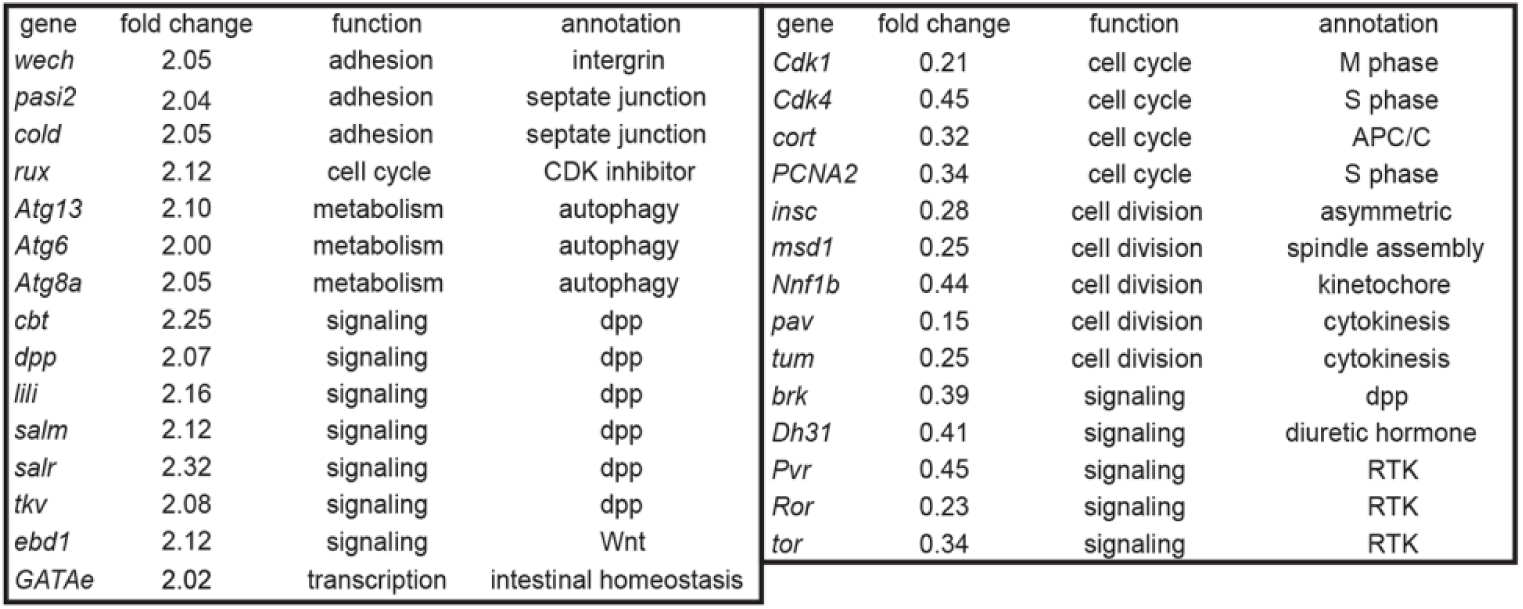
The T6SS promotes a unique transcriptional response from the intestine. Genes uniquely regulated in response to C6706 from RNA-seq of *Drosophila* whole guts.

**Supplemental Figure 4.**
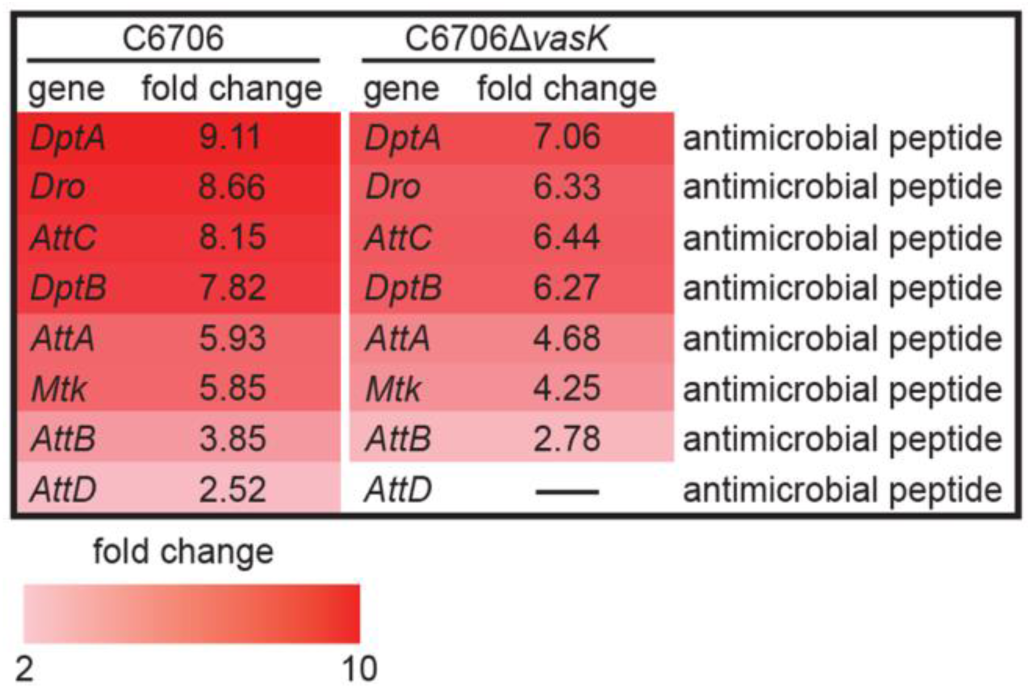
Infection with *V. cholerae* promotes the transcription of antimicrobial peptides. Antimicrobial peptide expressed from RNA-seq of *Drosophila* whole guts *infected* with C6706 or C6706Δ*vasK*.

## ACKNOWLEDGEMENTS

We are grateful for *CB>mCD8:: GFP* flies provided by Dr. Bruno Lemaitre, *esg[ts]>GFP* flies provided by Dr. Bruce Edgar, and *esg[ts]>CFP, Su(H)-GFP* flies provided by Dr. Lucy O’Brien. *Vibrio cholera* C6706 was provided by Dr. John Mekalanos and *Ecc15* was provided by Dr. Nicolas Buchon. We acknowledge microscopy support from Dr. Stephen Ogg and Greg Plummer at the University of Alberta. We acknowledge the FACS support provided by Dr. Aja Rieger at the University of Alberta. We acknowledge Kin Chan at the Network Biology Collaborative Centre (<u>nbcc.lunenfeld.ca</u>) for the IPC RNA-Seq service. This research was funded by grants from the Canadian Institutes of Health Research to EF (MOP77746) and to SP (MOP 137106). D.F, A.G, and B.K were supported by Nation Science and Engineering Research Council scholarships. M.F was supported by Alberta Innovates Technology Futures scholarship.

## DECLARATION OF INTEREST

The authors declare no competing interests

